# Plasma-free samples for transcriptomic analysis: a potential alternative to whole blood samples

**DOI:** 10.1101/2023.04.27.538178

**Authors:** Qingwang Chen, Xiaorou Guo, Haiyan Wang, Shanyue Sun, He Jiang, Peipei Zhang, Erfei Shang, Ruolan Zhang, Zehui Cao, Quanne Niu, Chao Zhang, Yaqing Liu, Yuanting Zheng, Ying Yu, Wanwan Hou, Leming Shi

**Author notes:** Corresponding authors: Ying Yu, Wanwan Hou Leming Shi. These authors contributed equally: Qingwang Chen and Xiaorou Guo.

## Abstract

RNA sequencing (RNAseq) technology has become increasingly important in precision medicine and clinical diagnostics and emerged as a powerful tool for identifying protein-coding genes, performing differential gene analysis, and inferring immune cell composition. Human peripheral blood samples are widely used for RNAseq, providing valuable insights into individual biomolecular information. Blood samples can be classified as whole blood (WB), plasma, serum, and remaining sediment samples, including plasma-free blood (PFB) and serum-free blood (SFB) samples. However, the feasibility of using PFB and SFB samples for transcriptome analysis remains unclear. In this study, we aimed to assess the viability of employing PFB or SFB samples as substitute RNA sources in transcriptomic analysis and performed a comparative analysis of WB, PFB, and SFB samples for different applications. Our results revealed that PFB samples exhibit greater similarity to WB samples in terms of protein-coding gene expression patterns, differential expression gene profiling, and immunological characterizations, suggesting that PFB can be a viable alternative for transcriptomic analysis. This contributes to the optimization of blood sample utilization and the advancement of precision medicine research.

## Introduction

RNA sequencing (RNAseq) technology (Wang et al. 2009; Stark et al. 2019) for transcriptomic analysis has become increasingly important in the fields of precision medicine (De Ruysscher et al. 2017) and clinical diagnostics (Ruan et al. 2021). RNAseq has also emerged as a powerful tool for identifying protein-coding genes (Wang et al. 2009; Hong et al. 2019), performing differential expression gene analysis for signature discovery (Soneson et al. 2015), and inferring immune cell composition (Newman et al. 2015). Human peripheral blood samples are widely used for RNAseq due to their availability and convenience of collection (Mohr and Liew 2007; Qi et al. 2023), offering valuable insights into the biomolecular information of the individual. As the transcriptome of blood samples has been more intensively studied (Hong et al. 2019), it has been shown that it can help improve the diagnosis of patients with rare diseases and identify new disease-related genes (Fresard et al. 2019). The analysis of blood transcriptome can provide prospective scientific guidance for disease prevention, diagnosis (de Almeida Chuffa et al. 2022), treatment (He et al. 2019a), and prognosis (Krishnan and Thomas 2022).

Blood samples can be classified into whole blood (WB), plasma, serum, blood cells, and other remaining sediment samples according to post-collection processing (Bayot and Tadi 2022). Plasma samples are obtained by centrifugation of WB samples with anticoagulants, while serum samples are obtained by centrifugation of WB samples without anticoagulants or with procoagulants (Sotelo-Orozco et al. 2021). WB, plasma, and serum samples are commonly used for transcriptomic studies due to their sample availability and convenience of collection under ideal conditions (Qin et al. 2016; Mjelle et al. 2019; Husseini et al. 2022). However, the remaining sediment samples after plasma or serum separation are often discarded or stored for biological resource preservation in real-world scenarios. These include plasma-free blood (PFB) and serum-free blood (SFB) samples, which also contain abundant RNAs that reflect individual information. WB samples contain various immune cell types, such as peripheral blood mononuclear cells (PBMCs), polymorphonuclear cells (granulocytes), and other anucleate cells (e.g., erythrocytes and platelets) according to the number of cell nuclei (Uhlen et al. 2019) (https://www.miltenyibiotec.com/US-en/resources/macs-handbook/human-cells-and-organs/human-cell-sources/blood-human.html). PFB samples are like WB samples but without plasma. Moreover, it should be noted that the cells in SFB samples are all enclosed in a blood clot, which may lead to differences in their expression when compared to WB and PFB samples.

For retrospective cohort studies that rely on biobank WB samples from authoritative institutes and hospitals for transcriptomic research, PFB and SFB samples may serve as alternative sources of RNA for sequencing when WB samples are insufficient. Nevertheless, the feasibility of using PFB and SFB samples for transcriptome analysis remains unclear. Moreover, as multiomics studies such as genomics, transcriptomics, proteomics, metabolomics, and microbiomics require adequate blood samples (Wu et al. 2021a), it is essential to maximize the utilization of blood samples and identify biomarkers in the blood for achieving personalized treatment (De Ruysscher et al. 2017; Kamali et al. 2022).

Given these concerns, our study aimed to assess the viability of employing either PFB or SFB samples as substitute RNA sources in transcriptomic analysis. Previous research have demonstrated the influence of pre-analytical factors on RNA quality and gene expression (Debey-Pascher et al. 2011; Mastrokolias et al. 2012; Dvinge et al. 2014; Shin et al. 2014; Zhao et al. 2014; Huang et al. 2017; Reust et al. 2018; Shen et al. 2018; Donohue et al. 2019; Gautam et al. 2019; He et al. 2019b; Harrington et al. 2020; Xing et al. 2021; Chebbo et al. 2022; Husseini et al. 2022). However, a comprehensive understanding of the similarities and differences among WB, PFB, and SFB samples in transcriptomic research remains elusive, making it crucial to evaluate their suitability for various applications using reliable metrics.

In the comparative study, we analyzed the transcriptomic profiles of WB, PFB, and SFB samples collected from two healthy donors at two different holding times. We assessed expression patterns, differential expression genes (DEGs), and immunological characterizations for each blood type. Our results indicate that PFB samples may be a potential alternative to WB samples for transcriptomic analysis, as they exhibit expression patterns and performance metrics are more closely correspond to those of WB samples. This would provide a valuable resource for optimizing blood sample utilization in research and clinical applications. Further research is needed to confirm the suitability of PFB and SFB samples for various specific applications.

## Methods

### Blood sample collection and preparation

Informed consents were obtained from two healthy donors (P10: male; P11: female) and followed the study protocol approved by the Ethics Committee of Fudan University. We collected 5 mL of blood from each donor using anticoagulation tubes with K2-EDTA (Shanghai Aoxiang Medical Technology Co., Ltd.) and procoagulation tubes (Jiangsu Yuli Medical Instruments Co., Ltd.). One group of blood samples was processed after holding 0.5 h (H0) and another group was processed after holding 6 h (H6) at room temperature to simulate the real turnaround times. PFB samples were obtained by centrifuging blood samples collected in anticoagulation tubes and separating the plasma supernatants, while SFB samples were obtained by centrifuging blood samples collected in procoagulation tubes and separating the serum supernatants. For each donor, three tubes of 150 µL of WB samples, three tubes of 100 µL of PFB samples, and one tube of 1.20 mL of SFB samples were used for RNA extraction, and the remaining samples were snap-frozen in liquid nitrogen and stored at −80℃ in the refrigerator. The separated plasma and serum samples in this study were used in another parallel study for miRNA analysis using small RNA sequencing.

### RNA extraction and cDNA library construction

RNA samples were extracted by QIAzol lysis reagents (QIAGEN) with QIAcube Connect (QIAGEN) according to the manufacturer’s manual. RNA concentration and integrity were measured by Qubit 3.0 Fluorometer (Thermo Fisher Scientific) and Agilent 4200 TapeStation (Agilent Technologies (China) Co., Ltd.), respectively. We normalized RNA concentrations according to the volume ratio of PFB vs. WB samples and SFB vs. WB samples to make them comparable among different blood samples. RNA samples with a total RNA amount ≥ 100 ng and RIN score ≥ 3 were qualified to construct Ribozero RNAseq libraries for sequencing (Li et al. 2014).

Globin and Ribosomal RNAs were depleted using Ribo-off Globin & rRNA Depletion Kit (H/M/R) (Vazyme #N408), and cDNA libraries were constructed using VAHTS® Universal V8 RNAseq Library Prep Kit for Illumina (Vazyme #NR605). The concentrations of cDNA libraries were measured using a Qubit 3.0 Fluorometer (Thermo Fisher Scientific). The distribution of cDNA fragment size was measured by Qsep 100 Advance (BiOptic).

### RNA sequencing and data preprocessing

cDNA libraries were sequenced on the NovaSeq 6000 sequencing platform (Illumina) to generate paired-end reads (150 bp). Then, we followed a previously published protocol (Chen et al. 2022) for processing and quality control of raw FASTQ reads. Briefly, adapter sequences were trimmed using fastp v0.19.6 (Chen et al. 2018) and read quality was assessed, while after trimming using FastQC v0.11.5 (https://www.bioinformatics.babraham.ac.uk/projects/fastqc/). Potential contamination from other species or junction primers was detected by extracting the first 10,000 reads from the clean FASTQ files using FastQ Screen v0.12.0 (Wingett and Andrews 2018). The mapping quality of 10% bam file per sample was calculated using Qualimap v2.0.0 (Okonechnikov et al. 2016) for efficiency and cost-effectiveness purposes. Reads were aligned to GRCh38_snp_tran genome reference and quantified using HISAT v2.1 (Kim et al. 2019), SAMtools v1.3.1 (Li et al. 2009), and StringTie v1.3.4 (Pertea et al. 2015) pipeline (Pertea et al. 2016) with Ensembl gene models (version: Homo_sapiens.GRCh38.93.gtf).

Gene expression was normalized by Fragments Per Kilobase of exon model per Million mapped fragments (FPKM) and Transcripts Per kilobase of exon model per Million mapped reads (TPM) to remove the effect of gene length and library size (Zhao et al. 2021). The counts were used for correlation analysis and to define detected genes for Jaccard index (Mukherjee et al. 2021) comparison, principal variance component analysis (PVCA), as well as DEGs analysis. FPKM was used for coefficient of variation (CV) calculation, sex check, principal component analysis (PCA), and hierarchical clustering analysis (HCA). TPM values were deconvoluted to infer the proportion of immune cell expression. Only protein-coding genes were used for CV calculation, PCA and correlation analysis. To avoid infinite values, a minimum value of 0.01 was added to the FPKM value of each gene before the log2 ratio.

### Sex-specific gene expression and sex check

Sex check was performed using sex-specific genes identified from previous studies (Chen et al. 2022), including five male-specific genes (*RPS4Y1*: Ribosomal Protein S4 Y-Linked 1, *DDX3Y*: DEAD-Box Helicase 3 Y-Linked, *EIF1AY*: Eukaryotic Translation Initiation Factor 1A Y-Linked, *KDM5D*: Lysine Demethylase 5D, *TXLNGY*: Taxilin Gamma Pseudogene, Y-Linked) and two female-specific genes (*XIST*: X Inactive Specific Transcript*, TSIX*: TSIX Transcript, XIST Antisense RNA). A distance matrix using the Euclidean method was calculated to measure the distance of the samples and genes, whereas Ward linkage was used for HCA using the R package pheatmap v1.0.12.

### Human protein-coding gene detection in different blood types

Similarities in protein-coding gene expression of all samples were compared using PCA. PCA was conducted with the univariance scaling, using the prcomp function of R package stats v4.2.1. Moreover, the clustering of protein-coding genes in the top 1000 standard deviation (SD) ranking across all samples was compared using HCA. HCA was performed using the R package pheatmap v1.0.12 with clustering based on a distance matrix calculated using the correlation distance metric. PVCA was performed using the R packages pvca v1.36.0 and Biobase v2.56.0 to assess the impact of different factors on the SD TOP 1000 protein-coding gene expression data. Pearson correlation coefficient (PCC) was calculated for the intra-group and inter-group to show correlations between technical replicates versus correlations between different blood types. The detected genes of the individual sample were defined as the genes with ≥ three counts in ≥ two technical replicates of three. The similarity of detected protein-coding gene sets between different groups was evaluated using the Jaccard index (Mukherjee et al. 2021). Finally, the detected protein-coding genes were filtered and compared between blood types with the same donor and the same holding time using R package eulerr v6.1.1 to produce Venn diagrams.

### Differential expression analysis and enrichment analysis

The R package limma v3.52.4 (Ritchie et al. 2015) was used to identify DEGs with a *p* < 0.05 and | log2FC | ≥ 1. Venn diagrams created by the R package eulerr v6.1.1 were used to compare and visualize the results. Gene Ontology (GO) and Kyoto Encyclopedia of Genes and Genomes (KEGG) enrichment analysis was performed using the R package ClusterProfiler 4.0 v4.4.4 (Roncaglia et al. 2017; Wu et al. 2021b) to assess the enrichment of DEGs in each group based on blood types or donors as variables.

### Immune cell cluster characterization and cell abundance

The CIBERSORT algorithm (Newman et al. 2015) was used to estimate the cell type composition of each sample using the R package IOBR v0.99.9 (Zeng et al. 2021), which translated the TPM-normalized gene expression matrix of different blood types into the relative proportion of immune cells. Euclidean distance was performed to assess the correlations between immune cell subsets using R package pheatmap v1.0.12.

### Immune cell-specific gene expression

The top 60 genes specifically expressed in six major cell types of human blood (granulocytes, monocytes, dendritic cells, NK-cells, B-cells, and T-cells) were collected from the "IMMUNE CELL" section of the website for Human Protein Atlas (HPA) (Uhlen et al. 2019), which can be accessed at https://www.proteinatlas.org/humanproteome/immune+cell. TPM values of these genes were used for cluster and comparative analysis in different blood types.

### Statistical analysis

All experiments were performed with three replicates. Statistical analysis and graphical work were performed using R v4.2.1 (https://cran.r-project.org/, R development core team) and a suite of R packages. Student’s *t*-test and Wilcoxon test were used to comparing RNAseq quality control metrics and gene expression profiles between groups with different blood holding times, donors, and blood types. All statistical tests were two-tailed, and a *p*-value less than 0.05 was considered statistically significant.

## Results

### Study overview

In this study, we performed comparative transcriptomic analysis of three blood types, including WB, PFB, and SFB, collected from two healthy donors at two different holding times (Fig. 1a, **top**). WB samples consist of diverse immune cell types, including PBMCs, polymorphonuclear cells (granulocytes), as well as anucleate cells (e.g., erythrocytes and platelets) based on their number of cell nuclei. PFB samples are similar to WB samples, while the cells in SFB samples are all enclosed in a blood clot (Fig. 1a, **bottom**). The RNA samples were subjected to library preparation and sequencing (Fig. 1b). The sequencing data were aligned to the hg38 reference genome, and transcript abundance was estimated for each sample. The quality check was performed using various metrics, including mapping ratio, genome region, CV, and sex check. To compare expression profiles across blood types, we employed various analytical methods such as PCA, HCA, PVCA, PCC, and Jaccard index. Additionally, we identified DEGs and compared immunological characterizations across the three blood types using different groups (Fig. 1c).

**Fig. 1.**
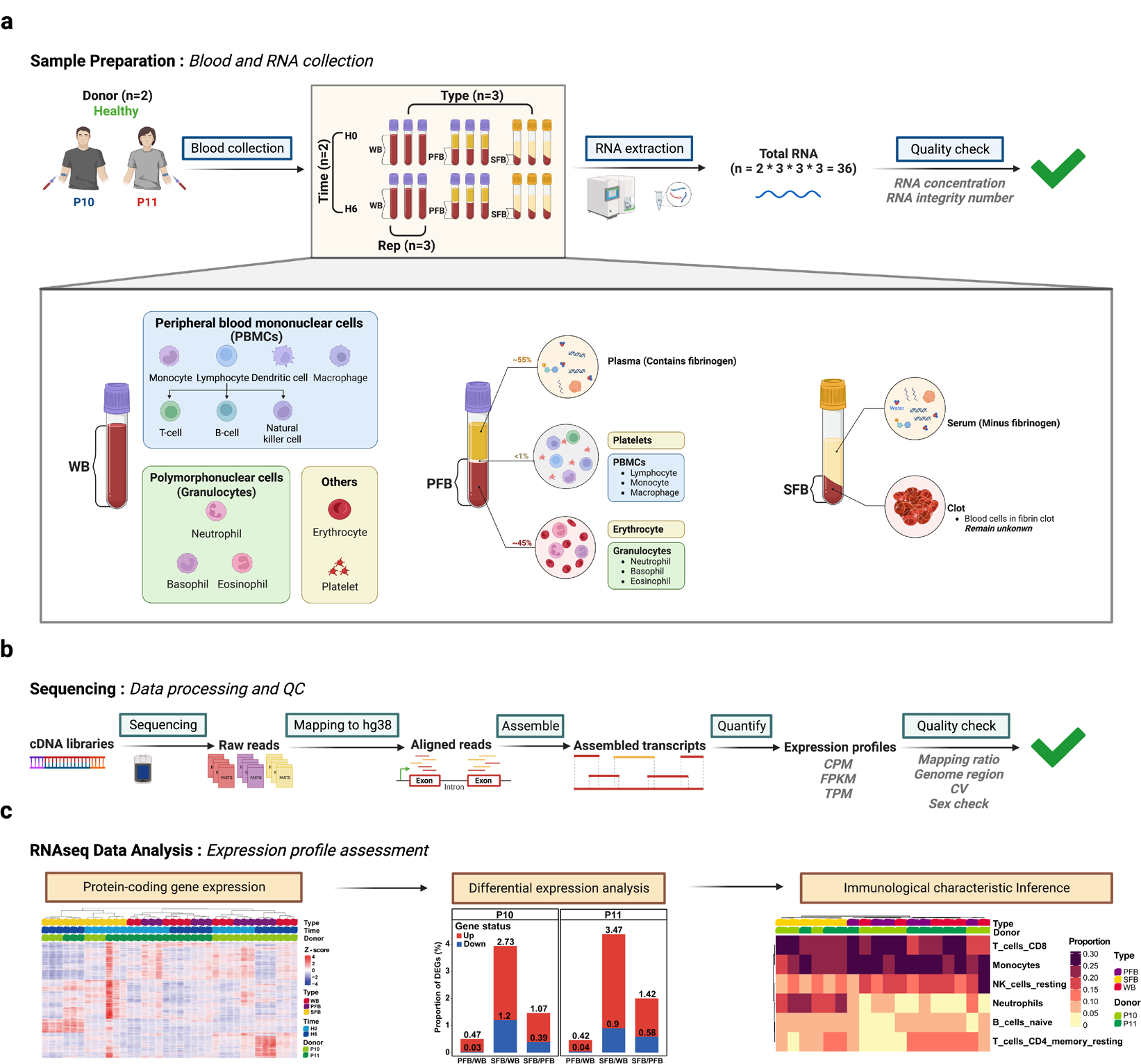
Overview of study design and analysis. **a Top**: Sample collection and processing. Whole blood (WB), plasma-free blood (PFB), and serum-free blood (SFB) samples were collected from two healthy donors (P10 and P11) through two holding times (H0 and H6). (P10: male donor, P11: female donor. H0: processed after holding 0.5 h, H6: processed after holding 6 h.) **Bottom**: Diagram of estimated immune cell composition in three blood sample types. Total RNA was extracted from each blood sample and assessed for quality, including RNA concentration and RNA integrity number (RIN). **b** RNA sequencing and quality control. The RNA samples were subjected to library preparation and sequencing. The sequencing data were aligned to the hg38 reference genome. Transcript abundance was estimated for each sample and the quality check was performed using various metrics, including mapping ratio, genome region, coefficient of variation, and sex check. **c** Bioinformatic analysis of RNAseq data. The expression of protein-coding genes across all samples was shown by HCA. Differential expression gene analysis was conducted between different blood types and donors. Immune cell composition was inferred for each blood type using gene expression data.

### High-quality RNA and sequencing data obtained from each blood sample

We assessed RNA quality by measuring RNA concentration and RIN score for each sample. The minimum RNA concentration required for cDNA library construction was 1 ng/μL (Li et al. 2014). All samples had RNA concentrations above this threshold with a median value of 34.50 ng/μL (Fig. 2a, **left;** Table 1). Furthermore, there was no significant difference in normalized RNA concentrations between the WB and SFB samples (**Fig. S1**). Similarly, the minimum RIN score required for sequencing was 3 (Li et al. 2014). All samples had RIN scores above this threshold, with a median value of 5.6 (Fig. 2a, **right;** Table 1). These results indicated that all samples were suitable for RiboZero library construction and sequencing.

**Fig. 2.**
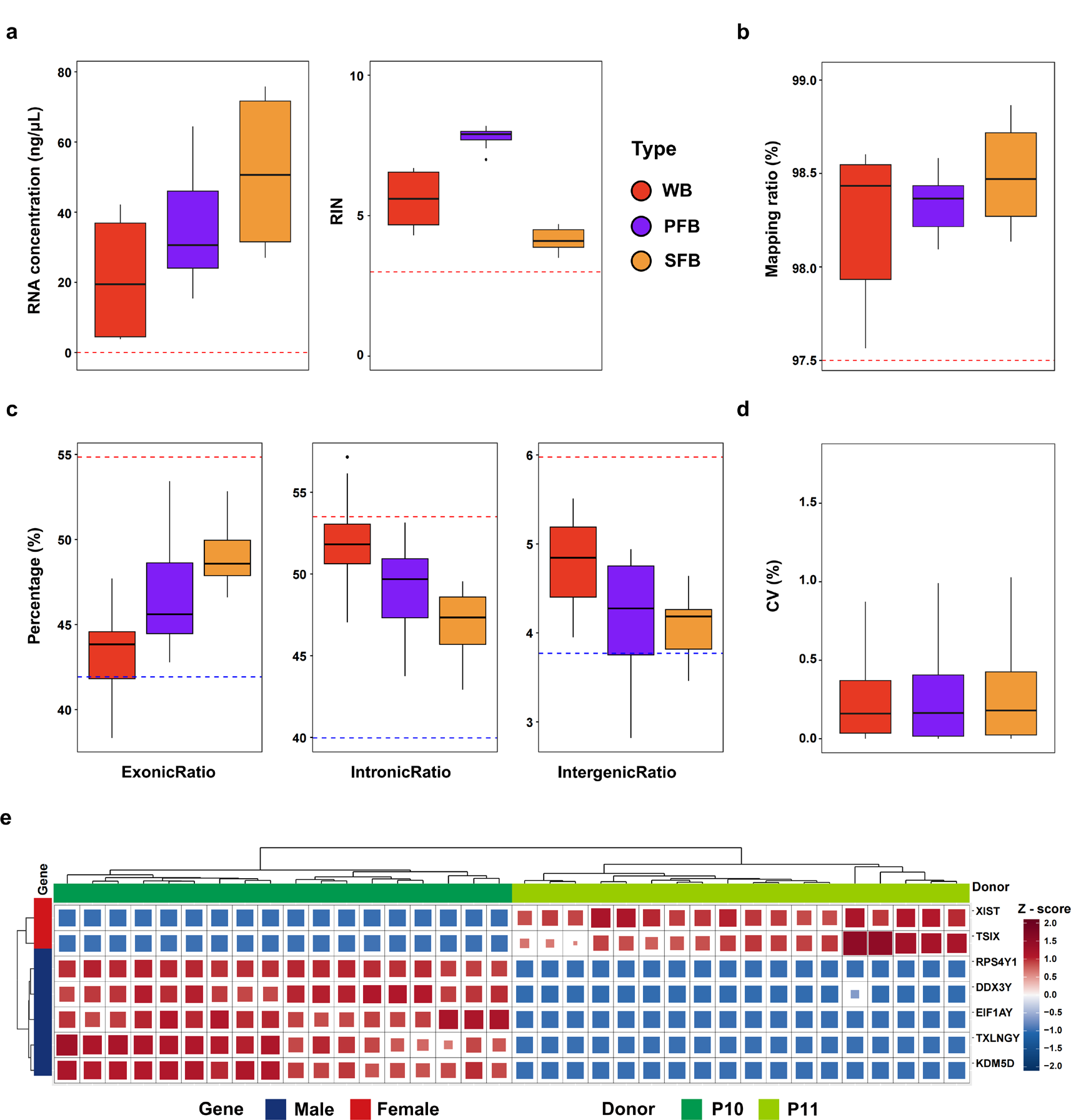
Quality assessment of RNAseq data. **a** RNA concentrations (ng/μL) and RIN values in three types of blood samples (Left red line value: 1; Right red line value: 3). **b** The mapping ratio (%) in three types of blood samples (The red line represents the median value (97.51%) corresponding to the RiboZero library mapping ratio in the Quartet dataset). **c** The distribution of reads across the human genome. The dotted line represents the mean ± sd values of the corresponding gene regions of the RiboZero library in the Quartet RNA reference dataset. Red indicates mean + sd value (ExonicRatio: 54.85, IntronicRatio: 53.51, and IntergenicRatio: 5.98), and blue indicates mean - sd value (ExonicRatio: 41.93, IntronicRatio: 39.97, and IntergenicRatio: 3.77). **d** The coefficient of variation (CV) values of protein-coding gene expression in three blood types **e** Heatmap and hierarchical clustering of the expression levels of seven sex-specific genes in all samples, including five male-specific genes (*RPS4Y1, DDX3Y, EIF1AY, KDM5D, TXLNGY*) and two female-specific genes (*XIST* and *TSIX*). Blue indicates male-specific genes, and red indicates female-specific genes. Seafoam green indicates the holding time H0, and dark blue indicates H6.

**Table 1.**
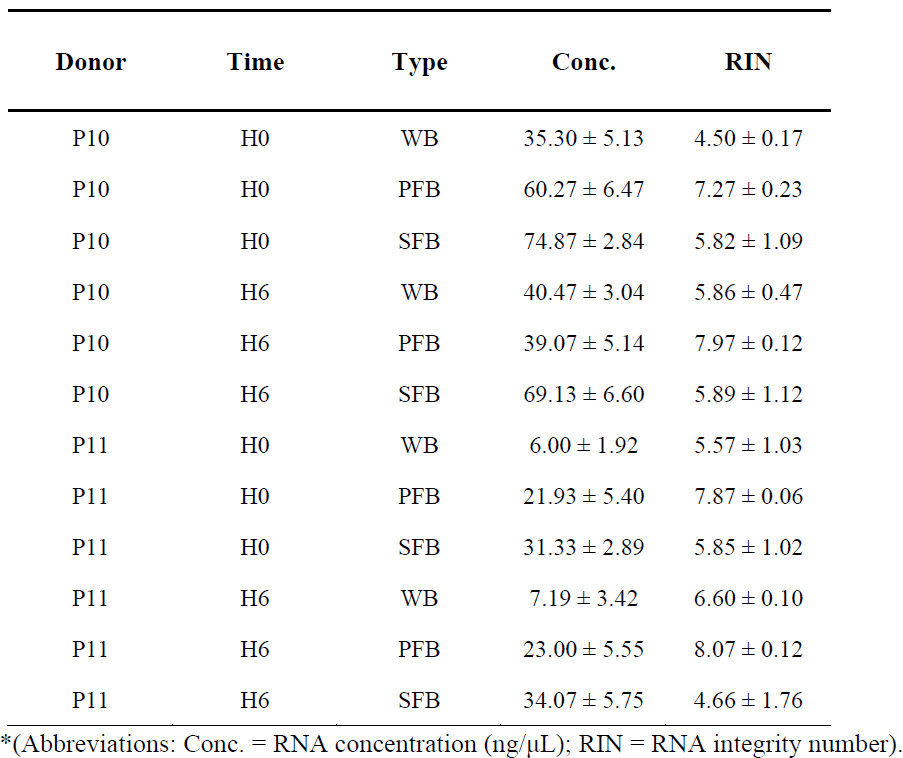
RNA Quantity and Quality *.

| Donor | Time | Type | Conc. | RIN |
| --- | --- | --- | --- | --- |
| P10 | H0 | WB | 35.30 ± 5.13 | 4.50 ± 0.17 |
| P10 | H0 | PFB | 60.27 ± 6.47 | 7.27 ± 0.23 |
| P10 | H0 | SFB | 74.87 ± 2.84 | 5.82 ± 1.09 |
| P10 | H6 | WB | 40.47 ± 3.04 | 5.86 ± 0.47 |
| P10 | H6 | PFB | 39.07 ± 5.14 | 7.97 ± 0.12 |
| P10 | H6 | SFB | 69.13 ± 6.60 | 5.89 ± 1.12 |
| P11 | H0 | WB | 6.00 ± 1.92 | 5.57 ± 1.03 |
| P11 | H0 | PFB | 21.93 ± 5.40 | 7.87 ± 0.06 |
| P11 | H0 | SFB | 31.33 ± 2.89 | 5.85 ± 1.02 |
| P11 | H6 | WB | 7.19 ± 3.42 | 6.60 ± 0.10 |
| P11 | H6 | PFB | 23.00 ± 5.55 | 8.07 ± 0.12 |
| P11 | H6 | SFB | 34.07 ± 5.75 | 4.66 ± 1.76 |

\*(Abbreviations: Conc. = RNA concentration (ng/μL); RIN = RNA integrity number).

The average mapping ratio of all samples was 98.36% (Fig. 2b), all above the median mapping ratio of 97.51% for the RiboZero libraries in the Quartet RNA reference datasets (Yu et al. 2022), which suggested the successful alignment of the reads to the reference genome. The gene region distribution was consistent with the expected characteristics of RiboZero libraries in the Quartet RNA reference datasets (Yu et al. 2022) across the three blood types (Fig. 2c). The median CV values of the sample replicates for the three blood types were all less than 0.17%, indicating good reproducibility of expression data (Fig. 2d). Additionally, a sex check was performed to validate the gender of the sample donors. Specifically, seven sex-specific genes were analyzed, including five male-specific genes and two female-specific genes. The results of the sex check were found to be consistent with the expected gender of each donor (Fig. 2e), thus affirming the quality of our expression data and providing confidence in the subsequent analysis.

Moreover, we performed additional quality control analysis on various metrics of the raw and mapped data (**Fig. S2**). Our findings revealed that: (1) high Phred quality scores were observed across all bases and positions in the FASTQ files (**Fig. S2a**); (2) no contamination from extraneous sources was detected in any of the samples (**Fig. S2b**); (3) the GC content exhibited a normal distribution in all samples (**Fig. S2c**). Details of the quality control metrics were provided in **Supplementary Table 1**. Overall, high-quality RNA and sequencing data could be obtained from each blood type, by employing rigorous quality control measures. These results established a strong foundation for further investigation into comparative gene expression profiling.

### Consistent expression of human protein-coding genes in PFB and WB samples

Due to the limited sequencing depth of our data, we primarily focused on the examination of highly expressed protein-coding genes as compared to non-coding genes in our expression analysis to ensure the reliability of our findings. We performed PCA of the protein-coding gene expression profiles to evaluate the effects of the donor, holding time, and blood type. The result showed that data were primarily separated by blood type on PC1 and by donor on PC2, while samples from the same donor and holding time clustered together by blood type (Fig. 3a). Notably, the PFB and WB samples of donor P10 from H0 were more closely related to each other than to other samples, and the same was true for the PFB and WB samples of donor P11, indicating a higher similarity between PFB and WB samples. To further investigate this similarity, we performed HCA on the SD TOP 1000 protein-coding genes in each sample, revealing that PFB and WB samples clustered together, whereas SFB samples clustered separately (Fig. 3b). Additionally, we employed PVCA to evaluate the contribution of different factors to transcriptional profile heterogeneity. The main factor was blood type (technical factor), followed by the donor (biological factor) and holding time (technical factor) (Fig. 3c).

**Fig. 3.**
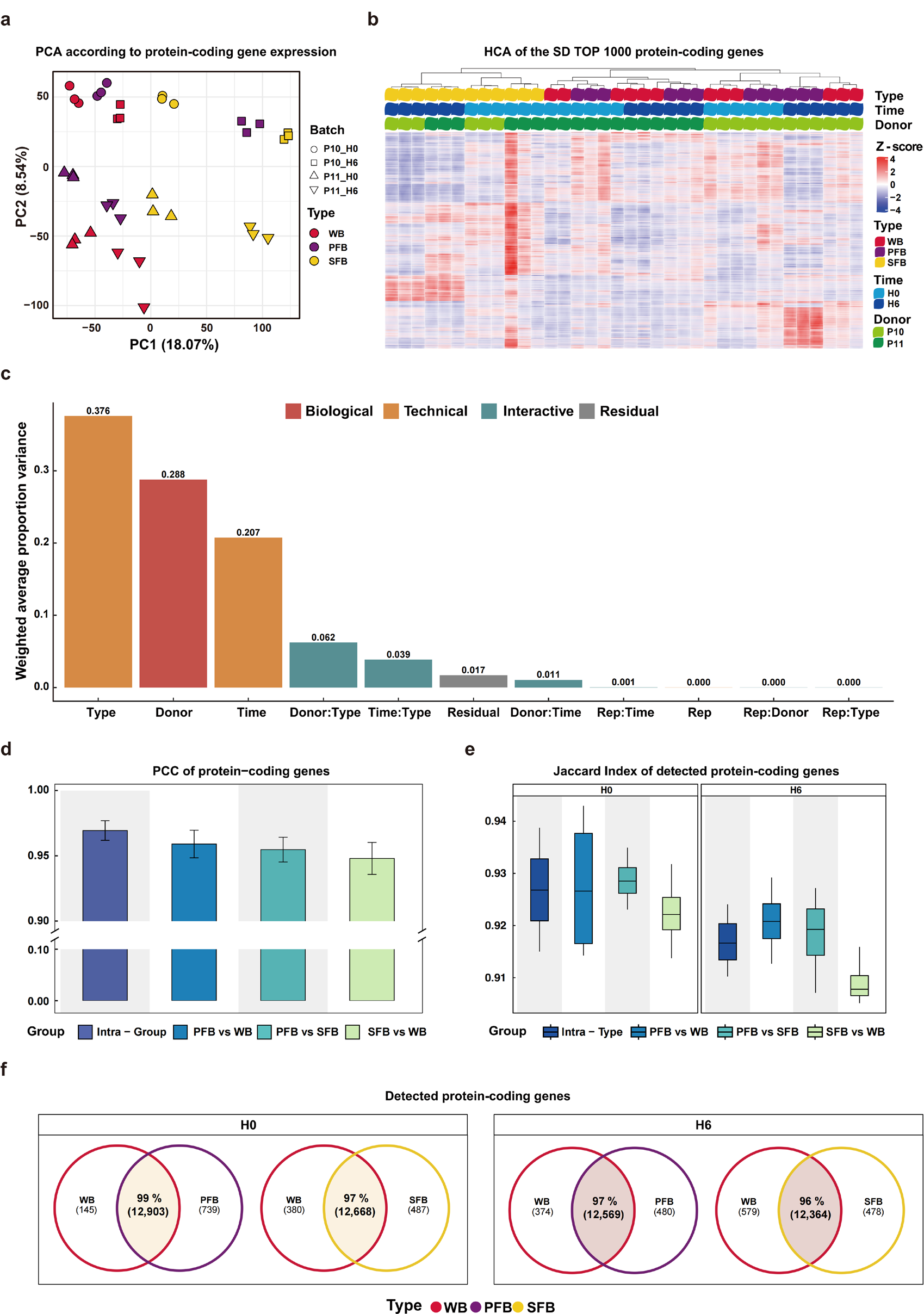
Human protein-coding genes expression analysis. **a** Principal Component Analysis (PCA) based on FPKM values of protein-coding genes in each sample (Red represents WB, purple represents PFB, and yellow represents SFB. The shape indicates the donor and the holding time of each sample.). **b** Heatmap of the top 1000 most variable (SD top 1000) protein-coding genes across all samples, selected and normalized by genes. "Type" represents the blood type, "Donor" represents the donor, and "Time" represents the holding time. **c** Principal Variance Component Analysis (PVCA) based on log2-transformed CPM values (+0.5) of the top 1000 SD protein-coding genes across all samples. **d** Barplot of the Pearson correlation coefficients for the protein-coding gene expression profiles (counts) in different sample groups, with the intra-group indicating the same blood type at different holding times. **e** Boxplot of Jaccard index of protein-coding genes expression levels (counts) in different groups at H0 and H6, with intra-type indicating samples from the same blood type in the same holding time. **f** Venn diagram of the number of detected expression genes in different blood types at H0 and H6.

Then, PCC was calculated for the protein-coding gene expression profiles (counts) in different sample groups. The expression levels of protein-coding genes were highly correlated across blood types, where the correlations between PFB and WB samples were closest to those between technical replicates (positive control: Intra-Group) (Fig. 3d). Detailed correlation analysis for all samples was presented in **Supplementary Figure 2**. Additionally, the Jaccard index was used to measure the consistency of gene detection among groups. Our analysis revealed that all groups had Jaccard index values above 0.91 (Fig. 3e) and that the Jaccard index values between PFB and WB samples were closest to those between technical replicates, indicating the similarity in protein-coding gene detection between PFB and WB samples. Last, we employed Venn diagrams to compare the number and proportion of detected protein-coding genes among WB, PFB, and SFB samples at H0 and H6. The results showed that over 96% of the protein-coding genes were detectable in PFB vs. WB samples and SFB vs. WB samples (Fig. 3f) at H0 and H6, demonstrating that each blood type was amenable to human protein-coding gene detection.

Above all, we demonstrated the consistent expression of human protein-coding genes in PFB and WB samples, highlighting their high similarity. However, to exclude the effect of holding time on our main findings (Fig. 3c), we only selected samples from H0 in subsequent analysis.

### Less variation between PFB and WB samples in differential expression analysis

To investigate the utility of the three blood types, we conducted a DEG analysis on WB, PFB, and SFB samples derived from two donors (P10 and P11) across three blood type comparisons (PFB vs. WB, SFB vs. WB, SFB vs. PFB). As expected, we found that the DEGs proportion between the three types of blood samples were low, ranging from a minimum of 0.46% (PFB vs. WB in P11) to a maximum of 4.37% (SFB vs. WB in P11) of all detected genes. Specifically, we observed that the smallest proportion of DEGs was observed between PFB and WB samples, accounting for approximately 0.5% (Fig. 4a), which further indicated that PFB and WB samples may have similar gene expression profiles.

**Fig. 4.**
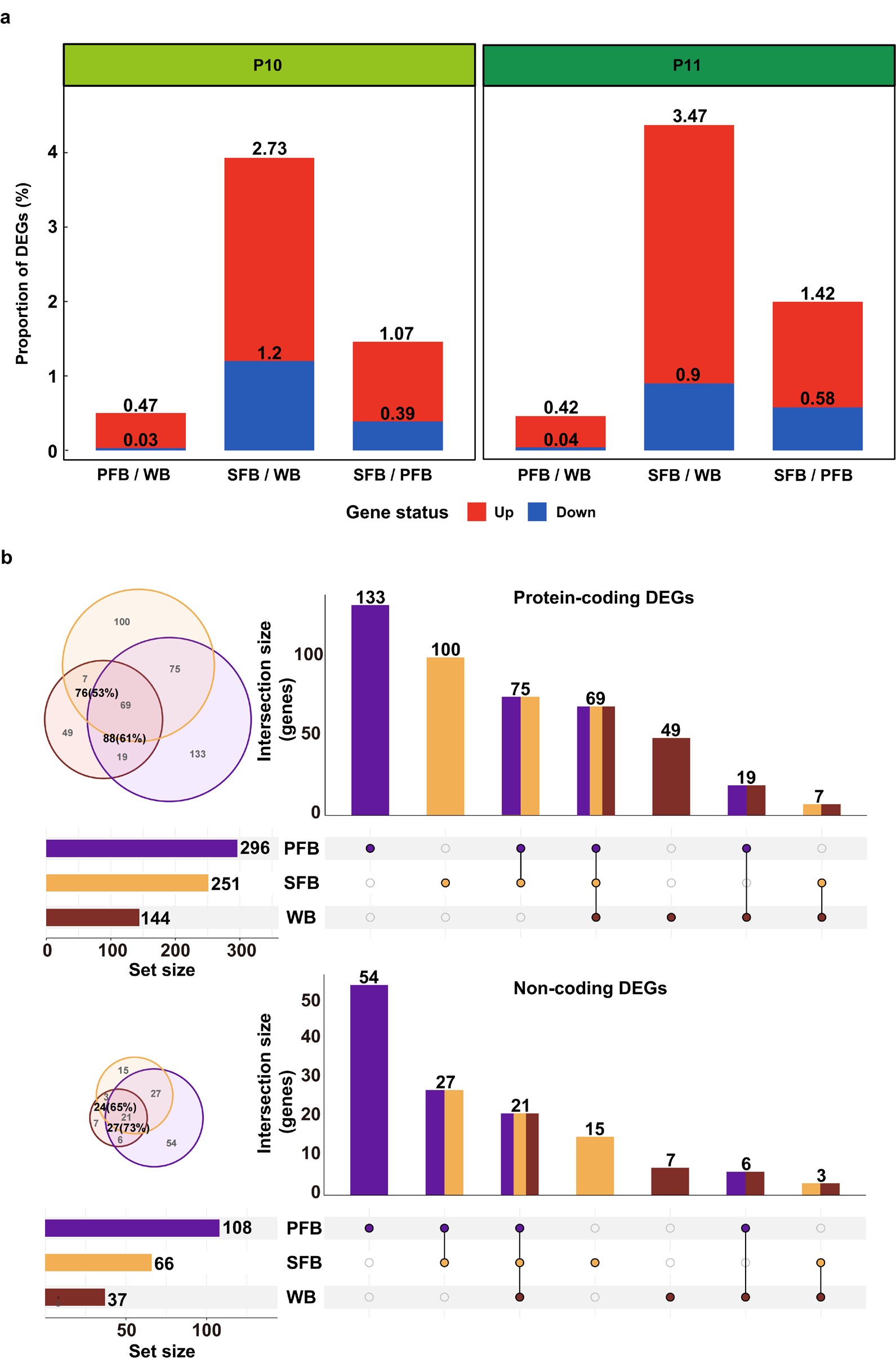
Differential expression genes analysis in different groups. **a** The percentage of differentially expressed genes (DEGs) between different blood samples from two donors. The comparisons are PFB vs. WB, SFB vs. WB, and SFB vs. PFB. Red indicates significantly up-regulated genes, and blue indicates significantly down-regulated genes. **b** The number of DEGs between two different donors for the three blood types. The comparisons are P10_WB vs. P11_WB, P10_PFB vs. P11_PFB, and P10_SFB vs. P11_SFB. The Venn diagram and upset plot provide an overview of the number of DEGs from the three blood types, including both protein-coding (up) and non-coding DEGs (down).

Moreover, we identified the DEGs between the two donors across three blood types (P10_WB vs. P11_WB, P10_PFB vs. P11_PFB, and P10_SFB vs. P11_SFB) to assess the ability of detecting biological difference. To determine the extent of overlap between DEGs, we compared PFB and SFB with WB samples. PFB samples covered 61% (88 in PFB / 144 in WB) DEGs of protein-coding genes and 73% (27 in PFB / 37 in WB) DEGs of non-coding genes in WB samples, while SFB samples can cover 53% (76 in SFB / 144 in WB) DEGs of protein-coding genes and 65% (24 in SFB / 37 in WB) DEGs of non-coding genes in WB samples (Fig. 4b). The results showed that PFB samples shared more DEGs with WB than SFB did, indicating that PFB samples were closer to WB samples in terms of differential gene expression analysis. Generally, a limited number of DEGs are observed between healthy individuals, and these are typically not utilized in pathway enrichment. However, the enrichment analysis of DEGs in WB samples revealed that only one pathway (external side of plasma membrane) was significantly enriched between donors, suggesting that inter-individual differences in gene expression may not have significant biological implications. These findings suggested that PFB samples exhibited a greater similarity to WB samples in terms of differential gene expression analysis compared to SFB samples.

### Similar properties of immune cell expression in PFB and WB samples

To compare the three blood types more comprehensively, we analyzed the composition of immune cells and the expression of related genes in WB, PFB, and SFB samples. The abundance of 22 types of infiltrating immune cells of each sample was estimated by CIBERSORT (Newman et al. 2015). We selected six cell types that had an average abundance of more than 1/22 across all samples for further analysis. HCA showed that WB and PFB samples clustered together while SFB samples formed a separate cluster (Fig. 5a). In addition, the inferred composition of immune cells in PFB samples showed greater similarity to that in WB samples, as compared to SFB samples, across the six immune cell types analyzed. The comparison between groups revealed that WB and PFB samples had no significant difference in immune cell composition except for neutrophils. SFB samples had higher proportions of neutrophils and a lower proportions of resting NK cells than other samples (Fig. 5b). This may be due to the influence of platelets and other components in WB, PFB, and SFB samples that affected neutrophils content and expression (Ruf and Ruggeri 2010).

**Fig. 5.**
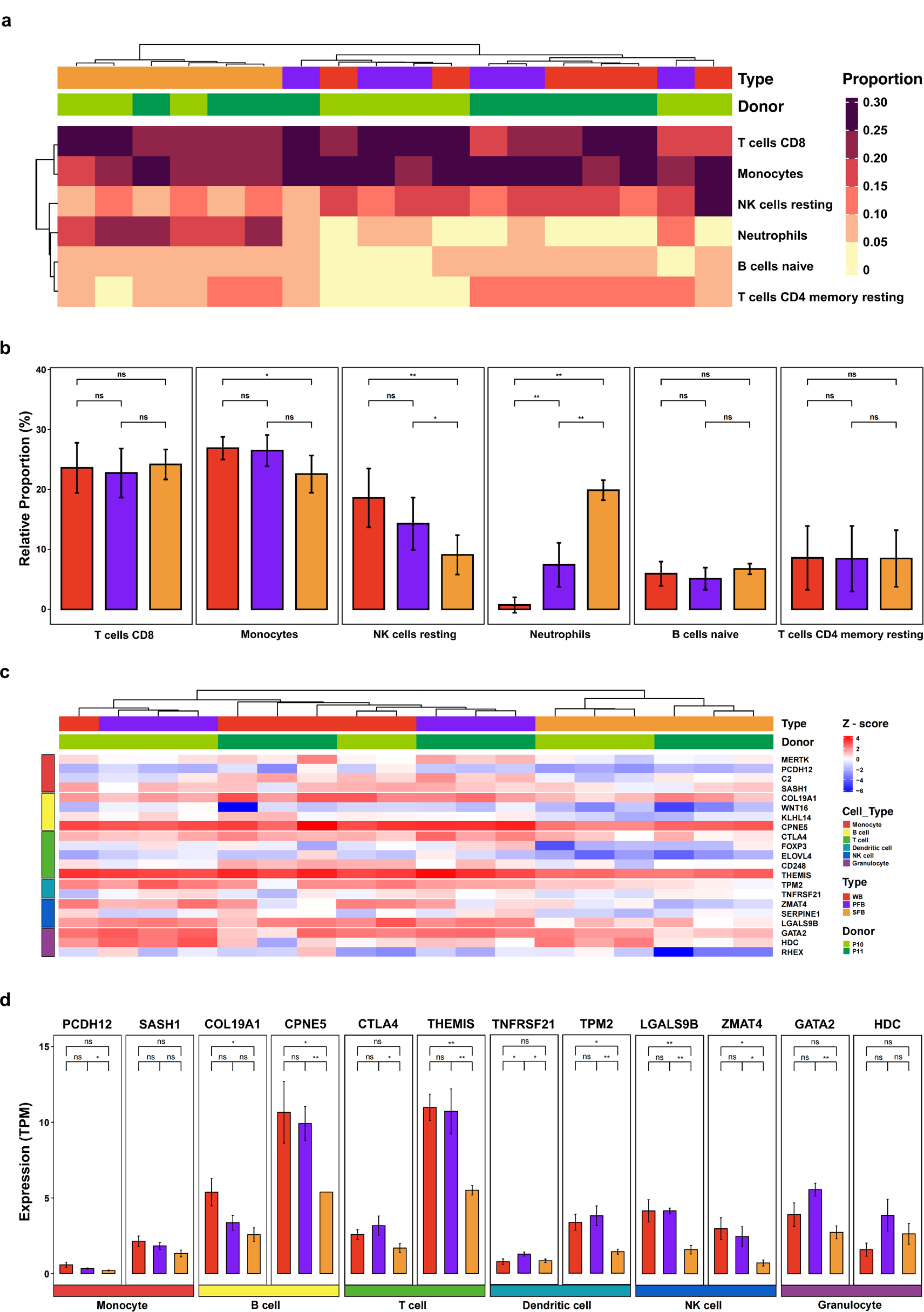
Inferred immunological characterizations. **a** Heatmap and hierarchical clustering of inferred relative abundance of immune cells in three blood types at H0 by CIBERSORT. Lime green indicates P10, and dark green indicates P11. **b** The proportion of six major immune cell types in each blood type at H0. **c** Heatmap and hierarchical clustering representing the expression levels of 21 immune cell-specific genes selected from the Human Protein Atlas (HPA) across six major immune cell types at H0. (Pale peach: Monocyte, deep pink: B_cell, magenta: T_cell, violet: Dendritic_cell, dark purple: NK cell, black: Granulocyte) **d** Comparison of expression levels of immune cell-specific genes between three blood types in six immune cell types.

Furthermore, we collected 60 immune cell-related genes specific to six major blood cell types from the HPA database. As different methods for classifying immune cells may vary, these six cell types of HPA may not entirely match the cell types from CIBERSORT results. After filtering out genes with unusually high or low expression levels across all samples, a subset of retained 21 genes was selected for subsequent clustering and comparative analysis. The clustering analysis produced consistent findings with the prior observations, signifying that WB and PFB samples also clustered together while SFB samples constituted a distinct cluster (Fig. 5c). Moreover, the comparative analysis between groups demonstrated that most immune cell-specific gene expressions were analogous between PFB and WB samples, whereas SFB samples exhibited certain discrepancies (Fig. 5d, **Fig. S2**). These results suggested that PFB and WB samples shared similar immunological characterizations.

## Discussion

Enhancing the utilization of blood samples can promote the advancement of precision medicine. In this study, we comprehensively compared transcriptomic profiles across three types of blood samples obtained from two healthy donors collected at two different holding times. The results implied that PFB samples may serve as a potential alternative to WB samples for specific applications where WB samples are impractical or unfeasible in retrospective and multiomics studies. We confined the study to two donors to maintain the reproducibility of inter-individual biological differences. Our findings exhibited a high correlation between PFB and WB samples at the expression profile level, with fewer DEGs between PFB and WB samples compared to SFB and WB samples. Notably, the individual difference (donor) profiles also demonstrated greater similarity between PFB and WB samples than between SFB and WB samples. Besides, we showed that PFB and WB samples had analogous gene expression traits of immune cell proportion and immune cell-specific gene sets. These observations bore substantial biological relevance, underscoring the feasibility of employing PFB samples as a practical substitute for WB samples in investigations about immune functionality and disease research (Feng et al. 2022). In summary, PFB samples could serve as an alternative to WB samples for RNAseq, thereby augmenting the significance of PFB samples in biobanks.

Interestingly, only one pathway (external side of the plasma membrane) was significantly enriched between donors in WB samples, indicating that the inter-individual differences in gene expression between two healthy individuals might not carry substantial biological implications. Additionally, discrepancies in neutrophil expression were observed between PFB and WB samples, suggesting that PFB samples might have certain limitations when employed as a surrogate for WB samples in the context of specific neutrophil-associated diseases (Wang et al. 2018; George et al. 2022; Gungabeesoon et al. 2023). Further investigation is required to clarify these differences and evaluate their potential consequences on the study of diseases about neutrophil function (Liew and Kubes 2019).

Apart from that, the availability of a broader range of blood samples for transcriptomics, including PFB samples, may expand the opportunities for investigating gene expression in diverse contexts. Several alternative blood types, such as resected clots from large vessel acute ischemic stroke (Tutino et al. 2021), dried blood spots (Reust et al. 2018), placenta and umbilical cord blood from pregnant women and newborns (Lu et al. 2022) have demonstrated promise in transcriptome analysis, yielding valuable insights into disease mechanisms and pathways. Consequently, the continued investigation of these alternative blood types is of paramount importance.

It is important to acknowledge the limitations of this study, which are as follows. First, the study utilized a relatively modest sample size, comprising blood samples from merely two healthy donors, each with three technical replicates. To bolster the validity of our findings, we advocated for the enlargement of the participant pool in subsequent research endeavors, incorporating multiple paired-donor comparative analysi<u>s</u>. Second, despite prior studies indicating that blood samples held for six hours did not significantly impact RNA concentration (Kim et al. 2007) and integrity (Huang et al. 2017), our RNA expression profiles showed a discernible degree of differences between inter-type samples and intra-type samples held for 0.5 h and 6 h. The causal mechanism of this observation remains unclear and needs further investigation. Last, as transcriptome analysis of blood samples predominantly involved protein-coding genes (Wang et al. 2020), our study mainly focused on protein-coding genes, and the expression patterns of non-coding genes in different blood types remains to be studied.

## Conclusion

This study confers a substantial understanding pertaining to the feasibility of deploying PFB samples as a viable substitute for WB samples in RNAseq applications. The findings present crucial perspectives and potential approaches for increasing the use of infrequently collected blood samples in biospecimen repositories for retrospective studies and maximizing the utilization of blood samples in multiomics cohort studies. Ultimately, the insights gleaned from this research contribute to the refinement of blood sample utilization in transcriptomic studies and the progression of precision medicine research.

## Abbreviations

RNAseq: RNA sequencing
WB: Whole blood
PFB: Plasma-free blood
SFB: Serum-free blood
PBMCs: Peripheral blood mononuclear cells
FPKM: Fragments Per Kilobase of exon model per Million mapped fragments
TPM: Transcripts Per kilobase of exon model per Million mapped reads
PVCA: Principal variance component analysis
DEGs: Differential expression genes
CV: Coefficient of variation
PCA: Principal component analysis
HCA: Hierarchical clustering analysis
RPS4Y1: Ribosomal Protein S4 Y-Linked 1
DDX3Y: DEAD-Box Helicase 3 Y-Linked
EIF1AY: Eukaryotic Translation Initiation Factor 1A Y-Linked
KDM5D: Lysine Demethylase 5D
TXLNGY: Taxilin Gamma Pseudogene, Y-Linked
XIST: X Inactive Specific Transcript
TSIX: TSIX Transcript, XIST Antisense RNA
SD: Standard deviation
PCC: Pearson correlation coefficient
GO: Gene Ontology
KEGG: Kyoto Encyclopedia of Genes and Genomes

## Acknowledgments

This study was supported in part by the National Natural Science Foundation of China (31720103909 and 32170657), the National Key R&D Project of China (2018YFE0201603, 2018YFE0201600, and 2021YFF1201305), Shanghai Municipal Science and Technology Major Project (2017SHZDZX01), State Key Laboratory of Genetic Engineering (SKLGE-2117), and the 111 Project (B13016). Fig. 1 was created with Biorender.com.

## Authors’ Contributions

L.S., Y.Z., and W.H. conceived the research and constructed the experimental design. L.S., Y.Z., Y.Y., and W. W. managed the project. Q.C., X.G., E.S., and C.Z. analyzed the data and participated in the verification and interpretation of data. W.H., H.W., S.S., P.Z., H.J., and Q.N. conducted blood sample collection, RNA purification, cDNA library construction, and quality control experiments. Y.L. and Y.Y provided some critical ideas for data analysis and manuscript. X.G. and Q.C. drafted the initial version of the manuscript. Z.C. and R.Z. contributed to the final revision of the paper. All authors have thoroughly reviewed and approved the final manuscript.

## Data Availability

The blood samples, RNA materials, and datasets generated during the current study are available from the corresponding author upon reasonable request.

## Code Availability

Our pipeline applications include all the Workflow Description Language (WDL, https://github.com/openwdl/wdl) processes required for RNAseq upstream analysis before the R analyses. Please note that the parameters in the pipeline applications are fixed, and the sample files can be directly processed. We have made the upstream analysis of RNAseq available on our GitHub repository (https://github.com/fudan-tnbc), and the dockers used in the upstream analysis can be obtained from D ocker Hub (https://hub.docker.com/u/chenqingwang). The code used for pre-proces sing the upstream result data and drawing the figures can be found on our GitHu b repository (https://github.com/xiaorouguo/c3bta). Quality control metrics for all data were collected in the metadata tables and visualized using R v4.2.1 (https://cran.r-project.org/, R development core team).

## Declarations

### Ethics approval

This study was approved by the ethics committee of the School of Life Sciences of Fudan University (Shanghai, China) (ID: BE2050).

### Consent to participate

Written informed consent was obtained from the participants.

### Consent for publication

All the participants approved to publish.

