## Supplemental files for "Plasma-free samples for transcriptomic analysis: a potential alternative to whole blood samples"

**Supplementary tables**

**Table S1. Metadata of RNA and cDNA library**

**Table S2. Metadata of RNAseq QC metrics**

**Supplementary figures**

**Fig S1. normalized RNA concentration**

**Fig S2. Quality control for clean data and sequence alignment**

**Fig S3. Pearson correlation coefficient analysis**

**Fig S4. Expression of 21 immune cell-specific genes**

**Supplementary Figure Legends**

**Fig S1. Normalized RNA concentration**

Normalized RNA concentrations (ng/μL) in the three types of blood samples.

**Fig S2. Quality control for clean data and sequence alignment**

**a** Line plot displaying high-quality scores across all positions in the FASTQ file for each sample. **b** Box plot illustrating mapping ratio (%) for all samples, differentiated by source (red represents whole blood (WB) samples, purple represents plasma-free blood (PFB) samples, and yellow represents serum-free blood (SFB) samples). **c** GC content for the three types of blood samples.

**Fig S3. Pearson correlation coefficient analysis**

Heatmap representing the Pearson correlation coefficients (PCC) for the protein-coding genes expression profiles (log_2_(counts + 1)) across all samples.

**Fig S4. Expression of 21 immune cell-specific genes**

Comparison of expression levels of 21 immune cell-specific genes among three blood types, spanning six immune cell categories.

**Supplementary Figures**

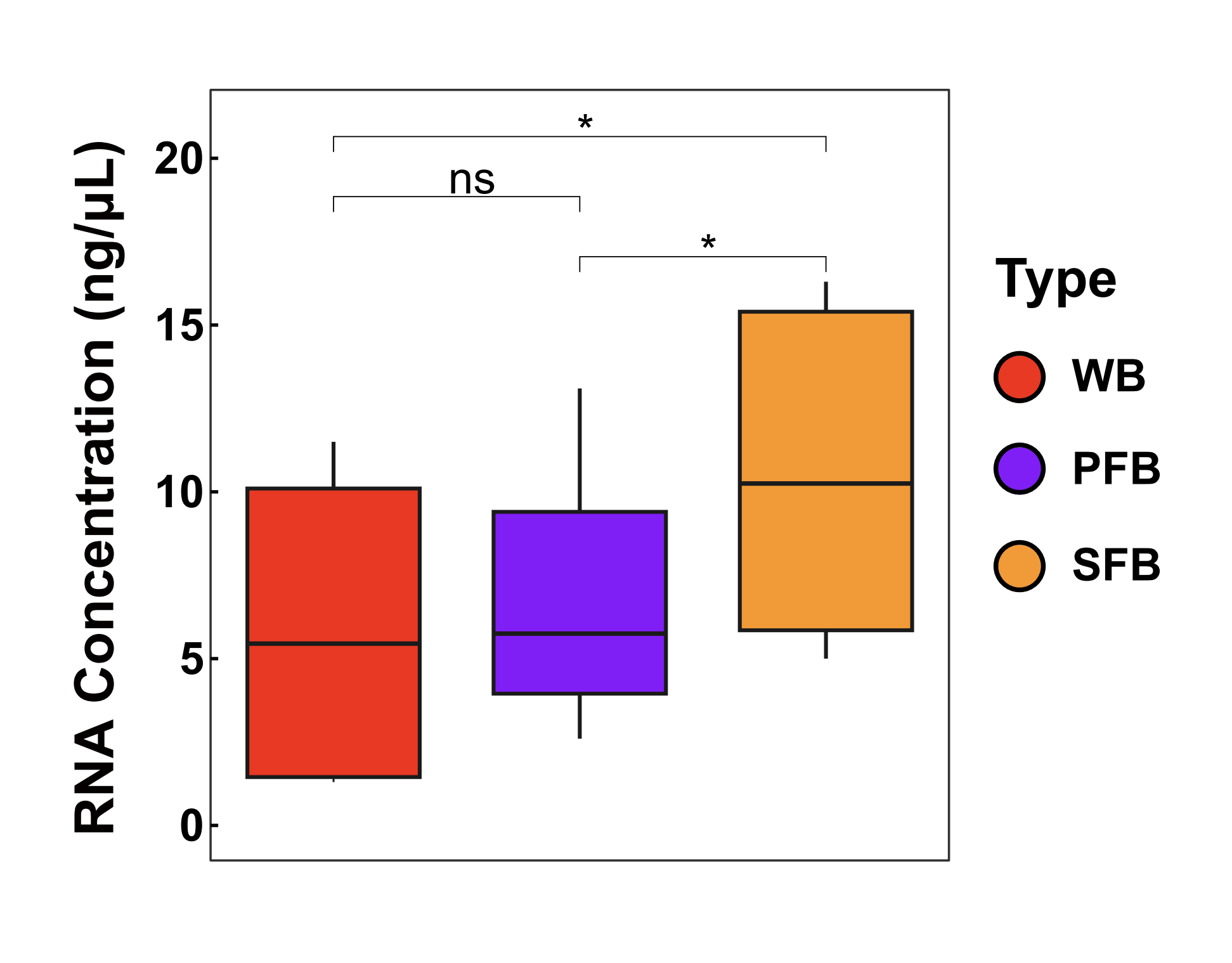

**Fig S1. Normalized RNA concentration**

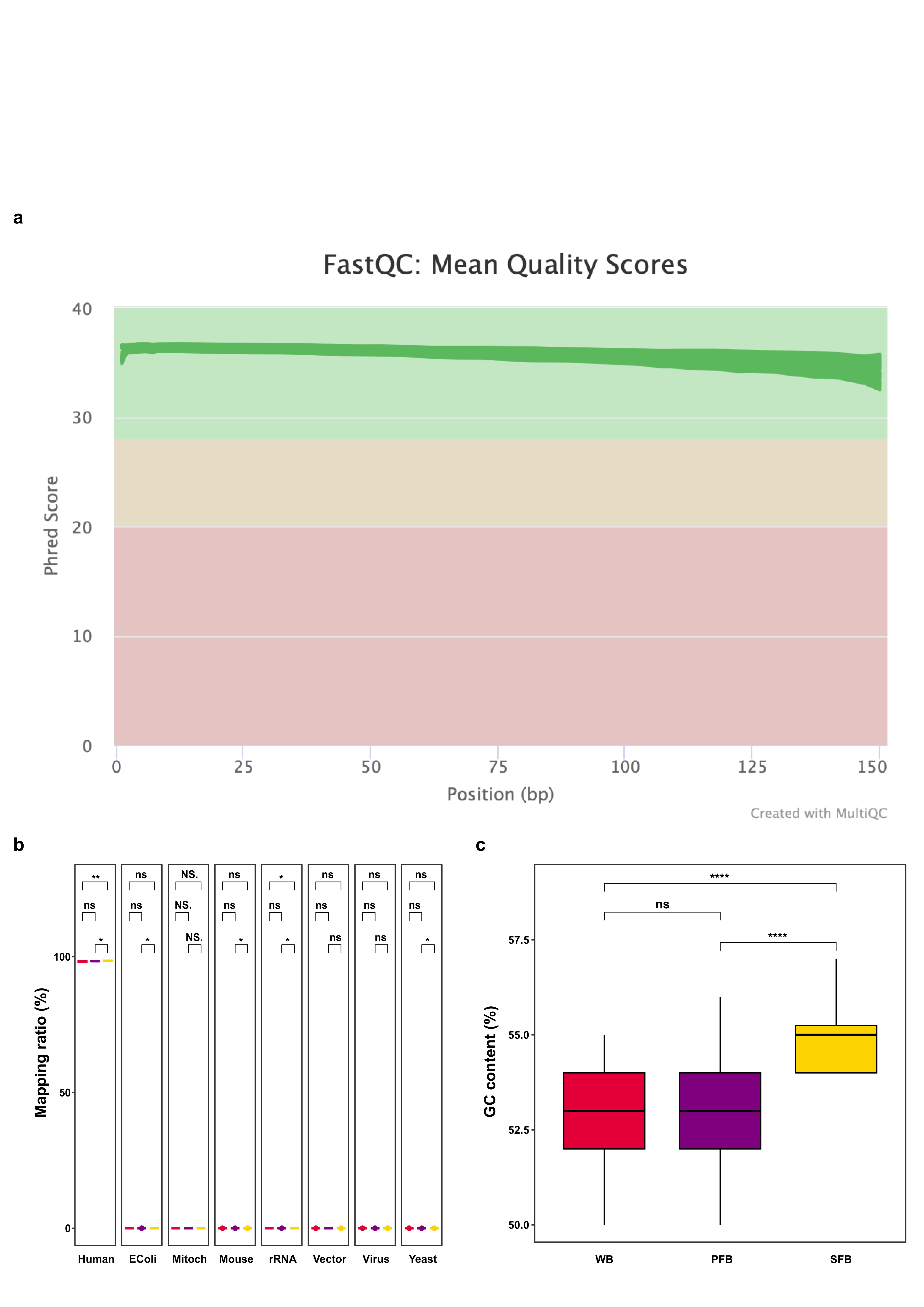

**Fig S2. Quality control for clean data and sequence alignment**

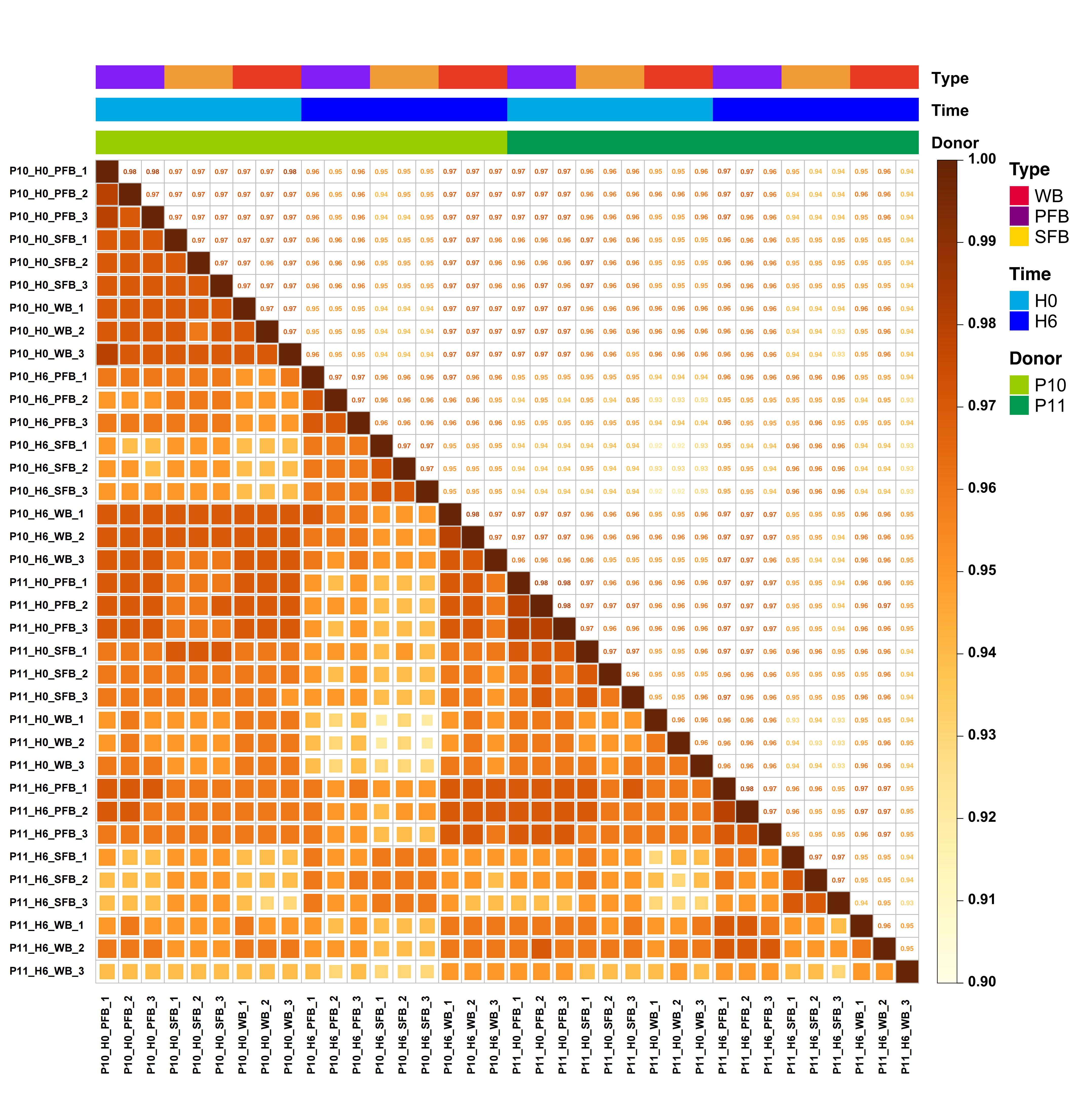
**Fig S3. Pearson correlation coefficient analysis**

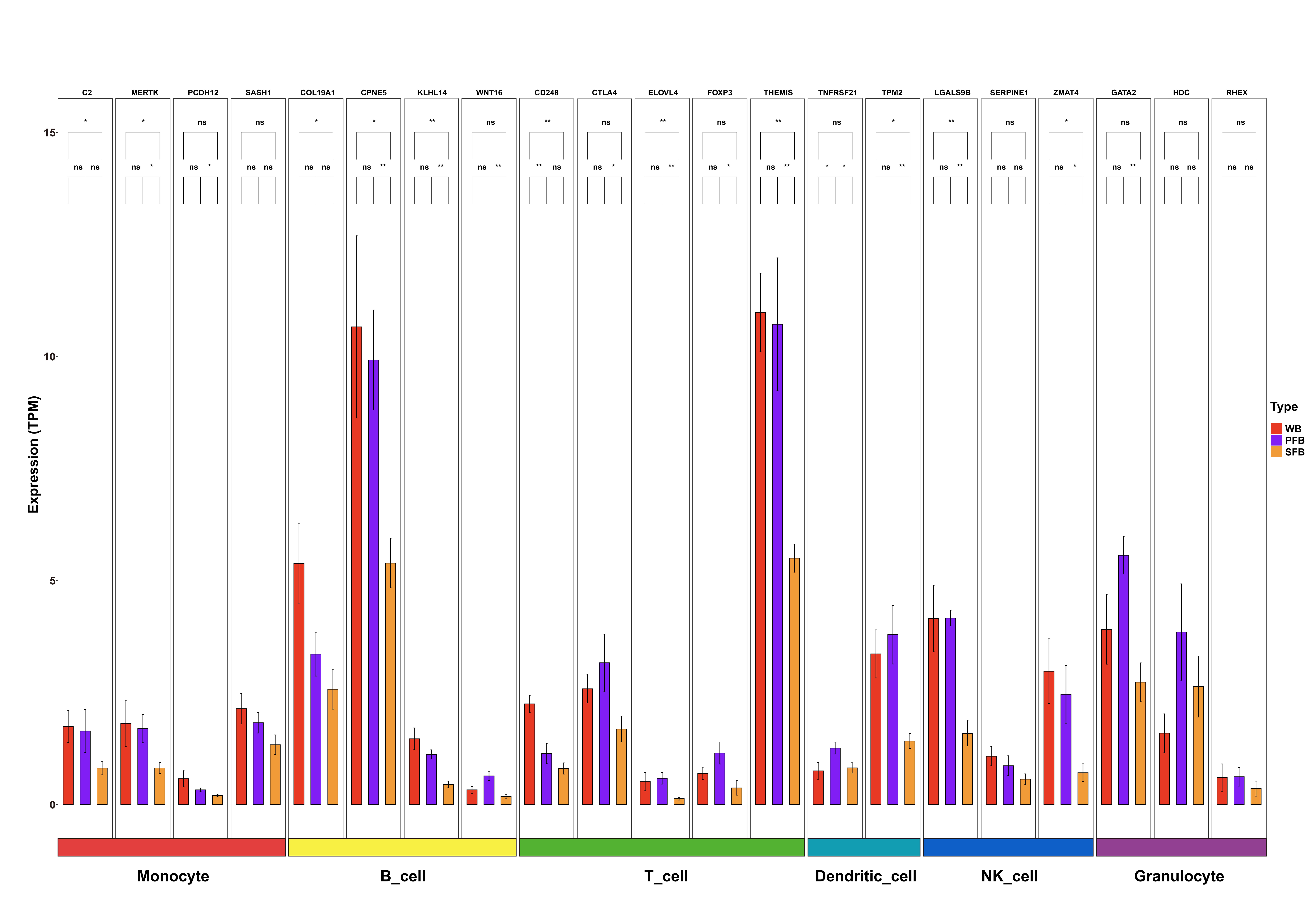

**Fig S4. Expression of 21 immune cell-specific genes**
